# Google searches accurately forecast RSV hospitalizations

**DOI:** 10.1101/607119

**Authors:** Benjamin M Althouse, Daniel M Weinberger, Samuel V Scarpino, Virginia E Pitzer, John W Ayers, Edward Wenger, Isaac Chun-Hai Fung, Mark Dredze, Hao Hu

**Author notes:** **Corresponding author:** Benjamin M Althouse, Institute for Disease Modeling, 3150 139th Ave SE, Bellevue, WA, 98005.

## Abstract

**Background:** Hospitalization of children with respiratory syncytial virus (RSV) is common and costly. Traditional sources of hospitalization data, useful for public health decision-makers and physicians to make decisions, are themselves costly to acquire and are subject to delays from gathering to publication. Here we use Google searches for RSV as a proxy for RSV hospitalizations.

**Methods:** Searches for “RSV” and numbers of RSV hospitalizations in WA, MD, FL, and CT were examined from 2004–2018. Running correlation coefficients and phase angles between search and hospitalizations were calculated. Various machine learning models were compared to assess the ability of searches to forecast hospitalizations. Using search data from all 50 US states, we use K-means clustering to identify RSV transmission clusters. We calculate the timing of the optimal timing of RSV prophylaxis initiation as the week beginning the 24-week period covering 95% of all RSV cases.

**Results:** High correlations (*>* 0.95) and low phase differences were seen between counts of hospitalizations and search volume in WA, MD, FL, and CT. Searching for RSV began in FL and radiated outward and three distinct transmission clusters were identified: the south and northeast, the northwest and Appalachia, and the center of the country. Calculated initiation dates for prophylaxis closely followed those calculated using traditional data sources (correlation = 0.84).

**Conclusions:** This work validates searches as a proxy for RSV hospitalizations. Search query surveillance of RSV is a rapid and no-cost addition to traditional RSV hospitalization surveillance and may be useful for medical and public health decision-making.

## Introduction

Clinicians, epidemiologists, and public health decision-makers now use search query surveillance, and other novel data streams, for timely reporting of incident cases of a pathogen [1] or for minimizing the effects of reporting bias for potentially sensitive topics [2]. Despite their recent surge in popularity, many novel data streams remain unvalidated. For example, does a Google search query for ‘flu symptoms’ or ‘depression symptoms’ indicate the searcher is suffering from influenza or depression themselves? Are they searching for general information, or for a loved one? This confusion around search intent led to errors in Google’s Flu Trends algorithm [3] that were later corrected by differentiating media-driven interest from actual infections [4, 5]. Understanding how these acts of information-seeking correspond to disease incidence, and which exact terms are used, is necessary if clinicians and public health officials are going to make reliable decisions based, at least in part, on data from novel sources.

Here, we focus on information-seeking surrounding respiratory syncytial virus (RSV), which causes significant morbidity and mortality among children under 5 years of age [6]. A leading cause of infant hospitalizations, RSV is responsible for over 100,000 childhood hospitalizations and $900 million to $4 billion in treatment costs annually in the United States [7]. Adult burden of RSV is also sizable – RSV has been identified in 2 to 5% of adult community-acquired pneumonias (CAP) [8]. Numerous vaccines are in development, with the three leading candidates focusing on maternal vaccination, with passive transmission of antibodies from mother to child, 2) direct vaccination of infants, and 3) vaccination of older individuals [9]. As of yet, the only licensed option for prevention of RSV is prophylactic administration of Palivizumab. Palivizumab is prohibitively expensive to administer to all children and thus is only used in high-risk infants, with monthly doses administered during the RSV season [10, 11]. Decisions to start a course of Palivizumab is made in relation to the prevalence of currently circulating RSV; thus, it is important to know rates of RSV disease in a population in near real-time. Additionally, theoretical work has suggested differential effectiveness of vaccine administration across the year for highly seasonal infectious diseases, with administration of a novel vaccine before the start of the oncoming season (when population level immunity is already high) having better outcomes than vaccination mid-season or at the end of the season [12].

Timely reporting of cases is important for situational awareness of RSV with which many additional public health decisions are made, such as adjusting hospital capacity [13, 14, 15] and clinically for understanding the potential etiology of CAP or increases in clinical severity in co-infections with other pathogens. Because effective clinical and public health decisions for RSV treatment rely on accurate, high-resolution situational awareness of outbreaks, a better understanding of the seasonal transmission patterns of RSV across geographical scales is needed. Using datasets containing every RSV-coded hospitalization in the states of Washington (WA), Maryland (MD), Florida (FL), and Connecticut (CT) since 2004 and Google Trends for Search, we validate searches for RSV as being indicative of RSV hospitalization. We then use searches for all 50 states to examine transmission patterns of RSV across the US.

## Methods

### Hospitalization data

Data on hospitalizations for RSV were obtained from the 2004-2015 State Inpatient Databases (SIDs) of the Healthcare Cost and Utilization Project (HCUP) maintained by the Agency for Healthcare Research and Quality (AHRQ) for WA, MD, and FL [16]. Hospitalization data from CT, which are in the same format as the HCUP data, were obtained from the CT State Inpatient Discharge Database through the CT Department of Public Health. All hospital discharge records from community hospitals in the states are included in the database. HCUP databases bring together the data collection efforts of State data organizations, hospital associations, private data organizations, and the Federal government to create a national information resource of encounter-level healthcare data [16]. We extracted all hospitalization records that included the International Classification of Diseases 9th revision, Clinical Modification (ICD-9-CM) code for RSV (079.6, 466.11, 480.1) listed as any one of the 25 listed discharge diagnoses. Data are reported at the monthly time scale for WA and CT, and quarterly time scale for MD and FL. The analysis of data from CT was approved by the Human Investigation Committees at Yale University and Connecticut Department of Public Health, Human Investigation Committee. The authors assumed responsibility for analysis and interpretation of these data.

### Search query data

Google searches for ‘RSV’ and ‘respiratory syncytial virus’ were monitored using Google Trends for all 50 US states and nationally from January 1, 2004 to May 31, 2018 and were normalized by the number of searches per 10,000,000 searches over the time period. We also monitored top Google search results for the term ‘RSV’ (searched June 8, 2018). Data were downloaded using the Trends Application Programming Interface for health.

### Normalizing data

Weekly search data were averaged into monthly (WA, CT) or quarterly (MD, FL) values, then a linear correction factor was applied to search data to scale to RSV hospitalizations. A simple linear regression was run with search as the predictor and hospitalizations as the outcome.

### Predicting hospitalizations

To assess the ability of search queries to predict hospitalizations – in advance of data releases by health departments and national agencies – we compared a model using search volume as normalized above to predict RSV hospitalizations to autoregressive integrated moving average (ARIMA) and negative binomial harmonic regression models, as well as an ensemble model. ARIMA models were fit using the algorithm by Hyndman and Khandakar’s [17], which automatically selects the best-fitting ARIMA model. Harmonic regressions were fit to the outcome of RSV hospitalizations using two harmonics (sine and cosine, annual and biennial periods) [18, 19]. We assessed the predictive ability of the models by leave-one-out cross-validation of the model fit. We focused the prediction on WA state RSV hospitalizations for parsimony. All analyses were conducted in R version 3.5.0 [20].

### Searches as surveillance

After validating searches in the four represented states, Fourier and wavelet transforms were taken on the log-transformed monthly time series of searches and phase angles of the time series calculated [21]. Based on Pitzer et al. [22], we examined the mean phase difference and phase correlations between all states and FL, as well as the correlation between the untransformed time series for each state. To assess the strength of biennial cycle in RSV searches we calculate the ratio of the biennial to annual Fourier amplitudes. To identify potential regions of correlated transmission we performed k-means clustering on the matrix of euclidean distances of search volume in all states and the phase angles for all states. We determined the number of clusters using the elbow method which adds clusters sequentially until adding more clusters does little to reduce the amount of variation between clusters [23, 24]. Finally, we calculated the optimal timing for administration of RSV prophylaxis as a rolling 24-week window from July to June of each year. We identified the window that has the largest proportion of RSV searches of the entire year (see Weinberger et al. [25] for details). Finally, we compare national level searches for RSV in the US and Australia.

## Results

Temporal trends in Google searches for ‘RSV’ per 10,000,000 were nearly identical to RSV hospitalizations over the study period, with high correlation between hospitalizations and searches ranging from 0.88 for Washington to 0.71 for CT (Figure 1), with small differences in absolute magnitude between numbers of searches and observed numbers of hospitalizations (Figure 1). Searches for ‘respiratory syncytial virus’ were of very low volume nationally (mean volume 8 searches per 10,000,000). Searches for ‘RSV’ in Florida tended to slightly precede hospitalizations. Running correlations indicate that the association between searches and hospitalizations is increasing over time. Unadjusted search volume remained above 0 in the summer months despite low numbers of RSV hospitalizations; this could be due to low levels of RSV circulation or searches unrelated to RSV incidence. Phase differences between searches and hospitalizations were nearly 0 across the study period (with FL searches slightly lagging hospitalizations), with phase differences trending towards 0 over time (Figure 2). In general, the prediction models performed well, with the models including both harmonic terms and searches being the best performing (Supplementary Table). Akaike information criterion, r-squared, and root mean squared error were all best for the combined model.

**Figure 1:**
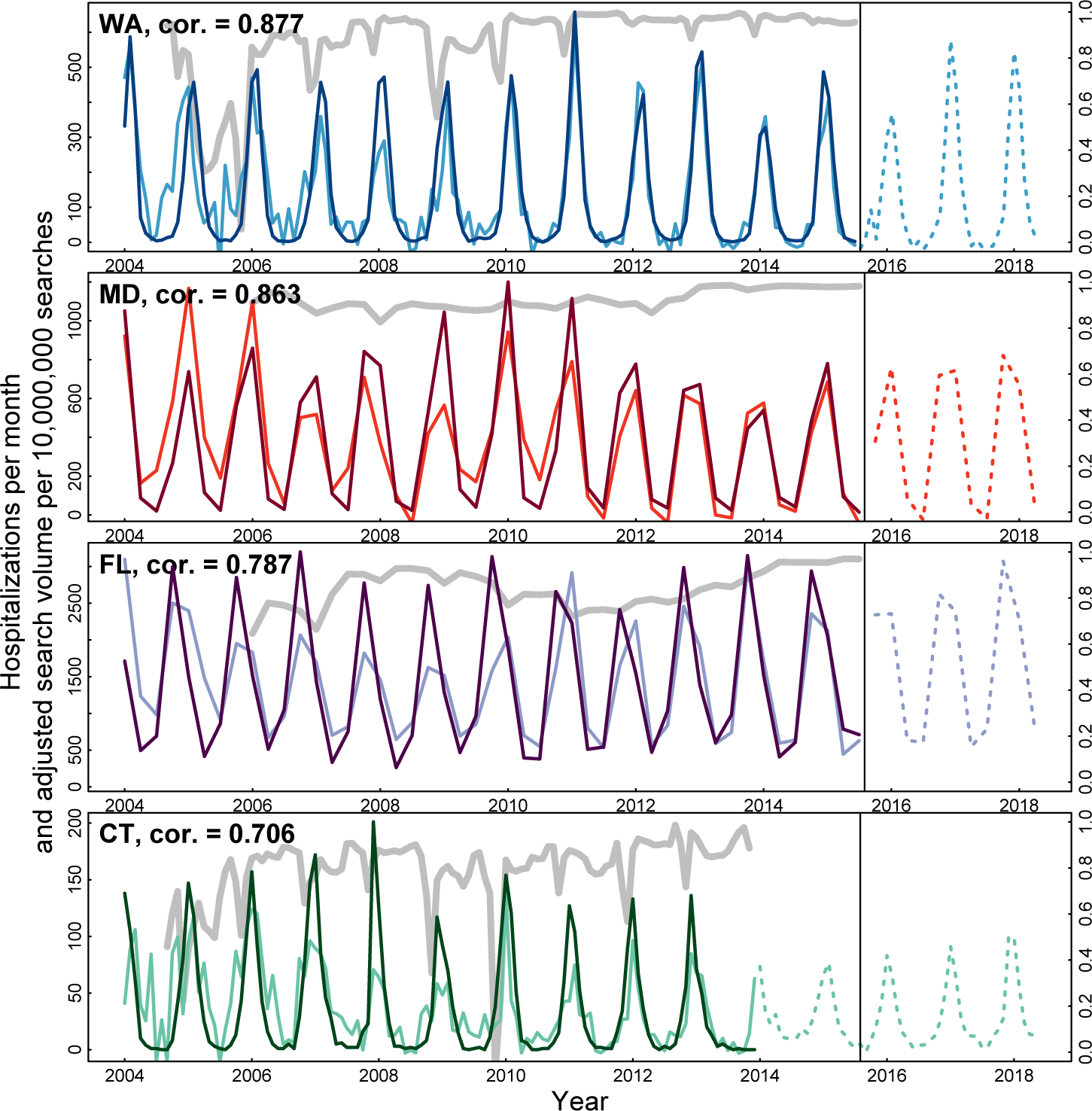
Summary of RSV search validation. Figure shows the numbers of RSV hospitalizations (dark lines) and search volume for ‘RSV’ (per 10,000,000 searches; light lines) for WA, MD, FL, and CT as well as predicted numbers of RSV hospitalizations for 2016, 2017, and 2018 RSV seasons. The grey lines are running correlations with a moving 9 month window.

**Figure 2:**
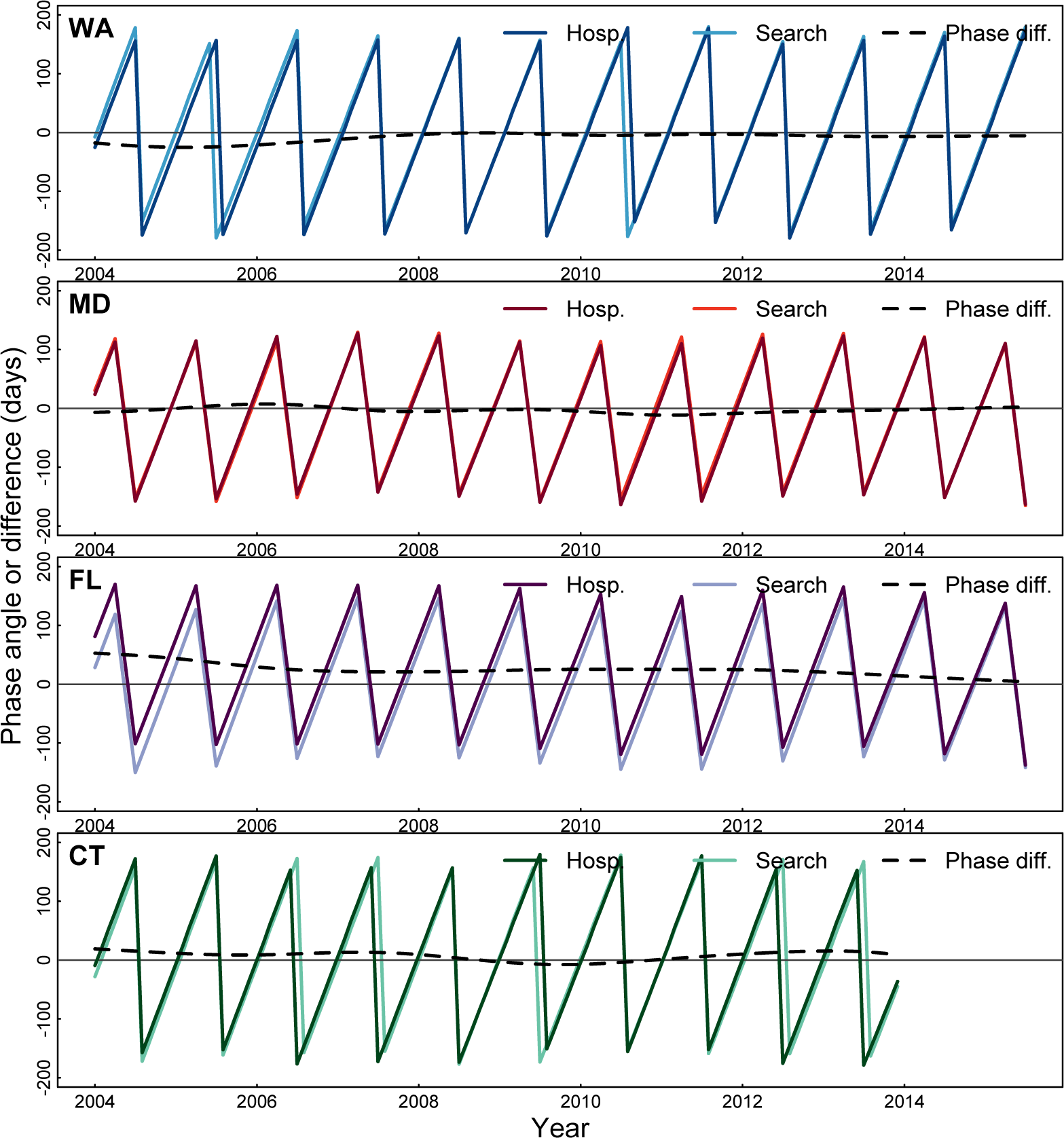
Phases of RSV hospitalizations and search volume. Figure shows the phase angles for hospitalizations and searches for WA, MD, FL, and CT and phase differences.

Using searches as a proxy for RSV activity reveals several notable patterns (Figure 3). First, annual dynamics begin in Florida and phase differences increase linearly with distance from Florida, with Montana and Oregon trailing Florida by nearly 50 days (Figure 3). Comparing the ratio of search volume in January and February to July and the ratio of Fourier 2-year to 1-year Fourier spectra shows the magnitude of seasonal change is the greatest in the Northern Rockies and middle of the country (Figure 4). Second, finding clusters of searches by through the k-means analysis of phase angles identifies 3 clusters of transmission (see Supplementary Figure 1): the south and northeast, the northwest and Appalachia, and a band across the middle of the US (Figure 3). This pattern is different when performing the k-means analysis on adjusted searches, with the west and east coasts in a cluster, the midwest in a cluster, and the Dakotas and Wyoming. Finally, we calculate the optimal timing for RSV prophylaxis by state (Figure 5) and find timings ranging from 15 weeks after July 1 for FL (October 12), to over 21 weeks past July 1 for WY (November 27). Finally, comparing northern and southern hemispheres, we see opposite patterns of RSV searches in Australia compared to the US (Supplementary Figure 2).

**Figure 3:**
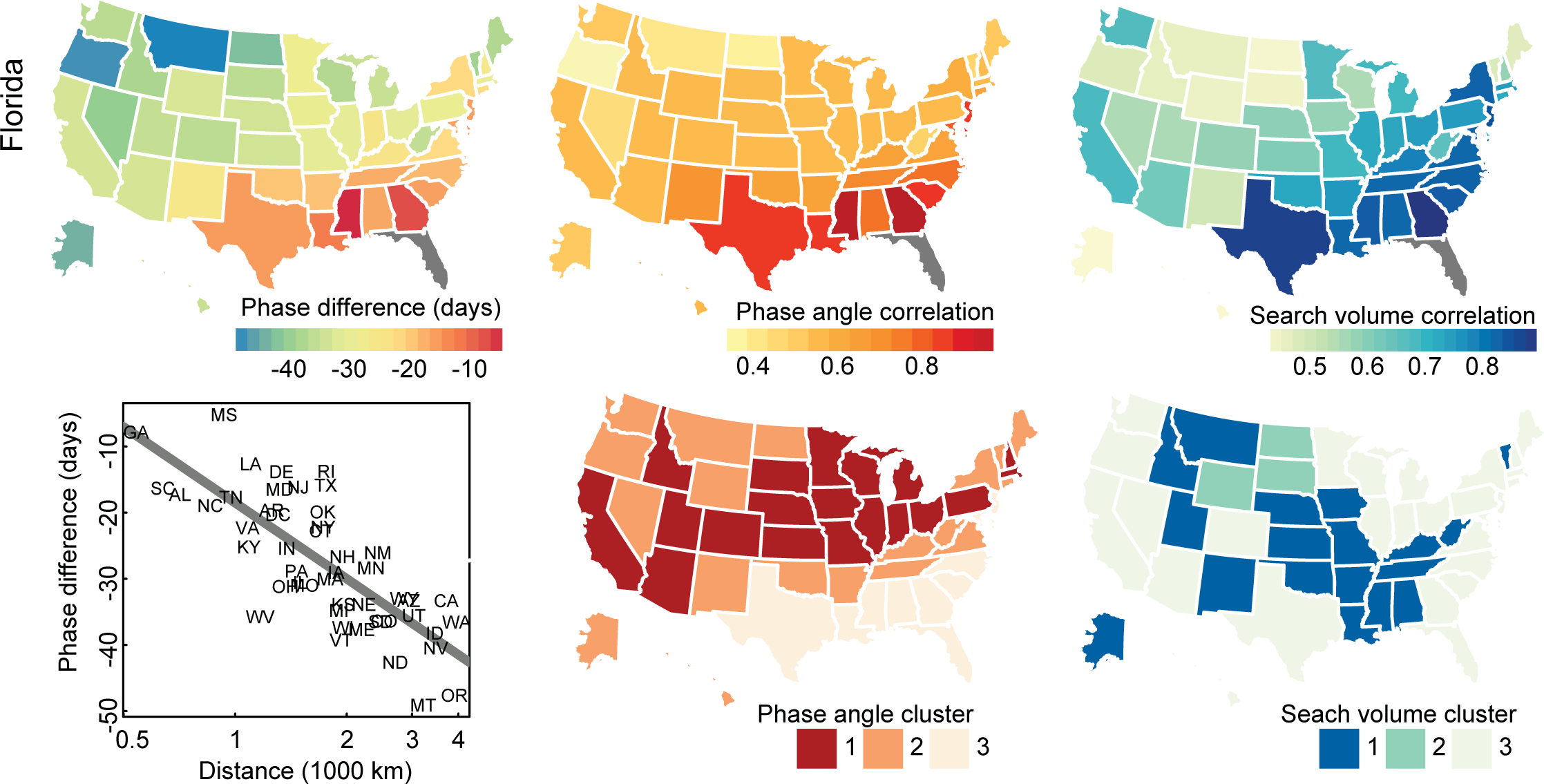
RSV searches as RSV surveillance. Figure shows the phase angles for RSV searches (left column), and correlations between phase angles (middle column) and raw searches (right column) relative to FL. Bottom left panel shows the phase difference as a function of distance from FL, where the line is a linear regression. The bottom right panels show clusters identified by k-means on euclidean distances between phase angles (bottom left map) and search volume (bottom right map).

**Figure 4:**
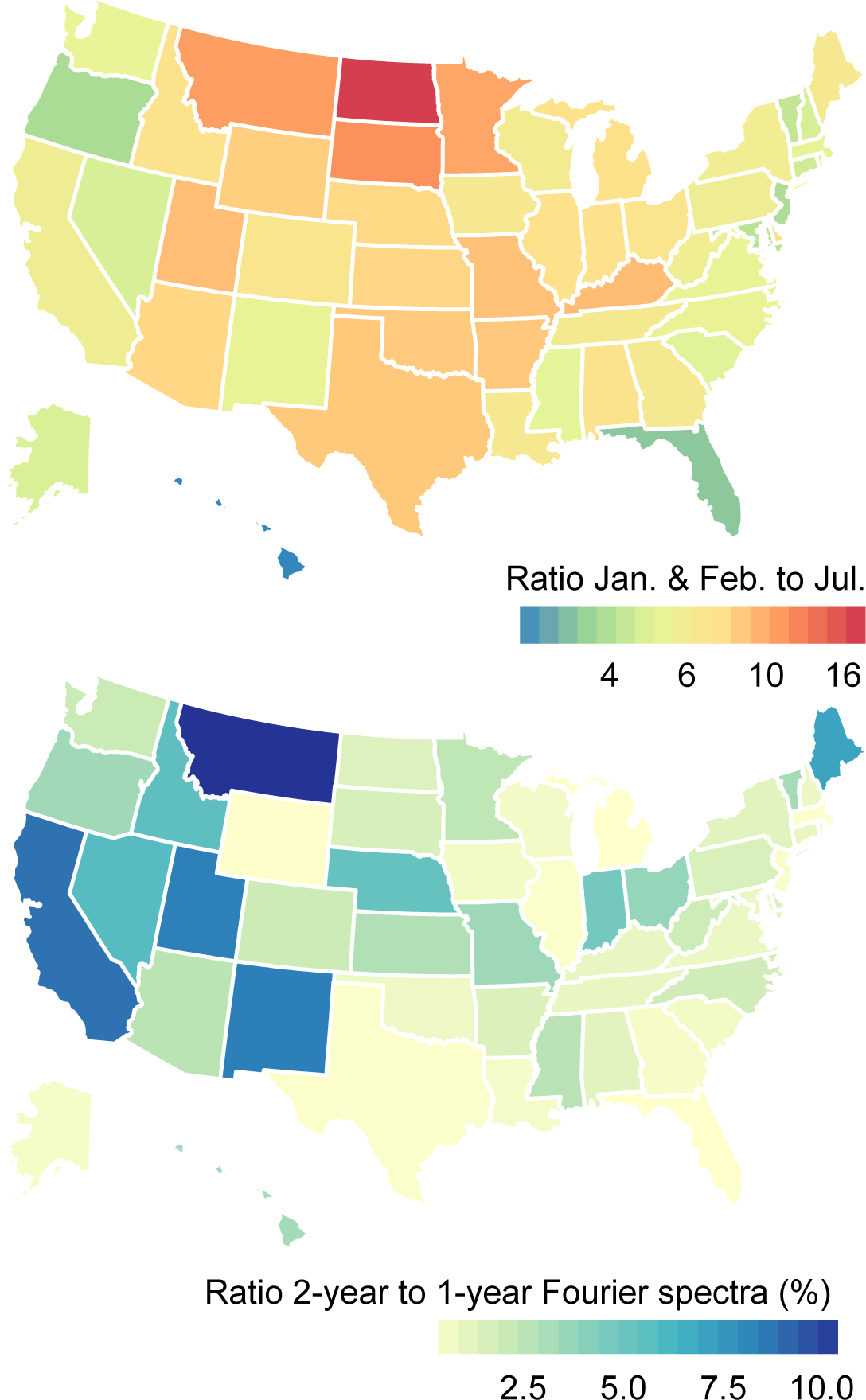
Ratio of winter to summer months and 2-year to 1-year Fourier spectra. Figure shows the ratio of search volume in January and February to July across the US (top map) and the ratio of the 2-year to 1-year Fourier spectra (bottom map).

**Figure 5:**
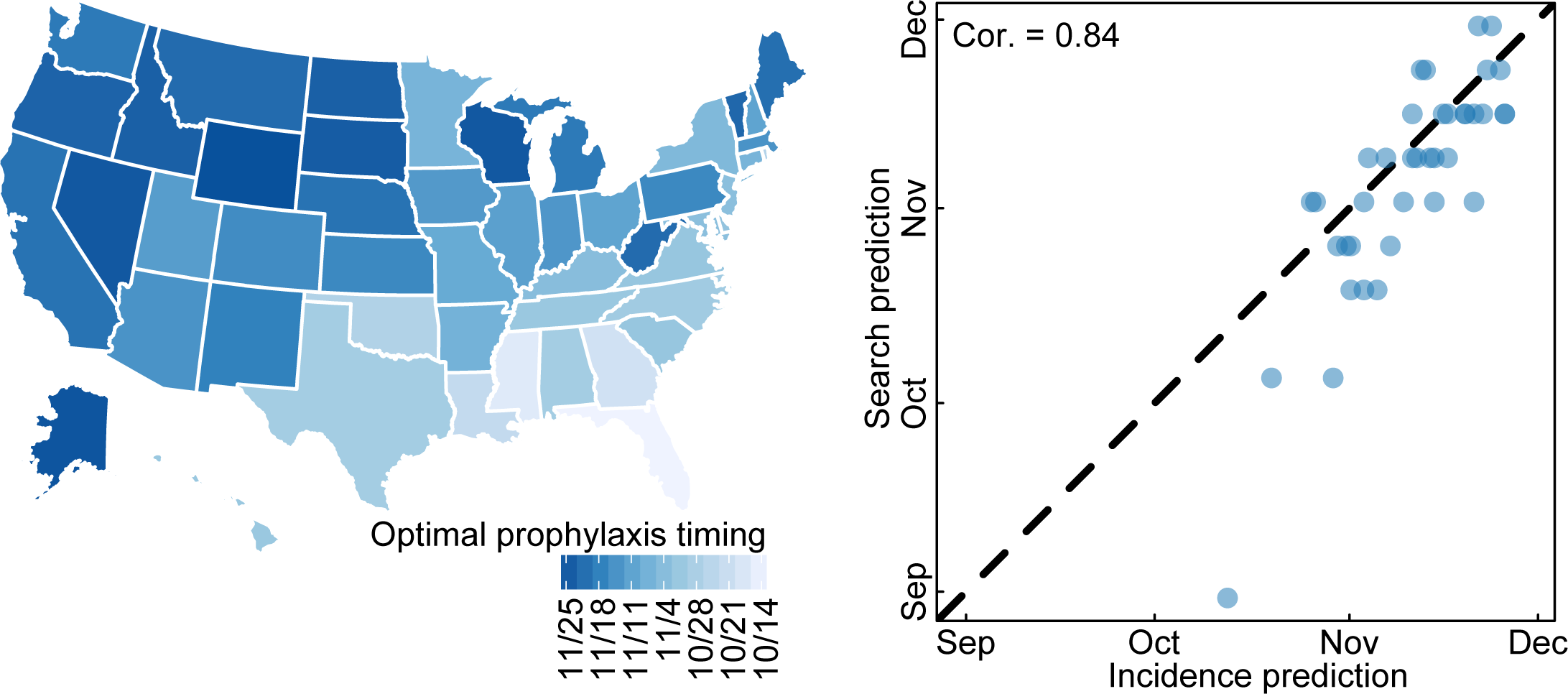
Optimal timing for RSV prophylaxis based on search volume. Figure shows the optimal numbers of weeks from July 1 for the start of prophylactic RSV treatment (left panel). Optimal numbers of weeks are averaged over the RSV seasons 2008-2009 to 2017-2018. The right panel compares the search-predicted optimal week with the weeks calculated in Weinberger et al [25].

## Discussion

Here we demonstrate the direct relation between a search query and a medical outcome, validating the use of searches for RSV in four states: WA, MD, FL, and CT. While the association between searches and outcomes should be validated in other localities – which may be problematic due to the granularity of reported hospitalizations (e.g., yearly or quarterly, as opposed to weekly or monthly) – the results presented here indicate a clear association between RSV diagnosis and online information-seeking behavior. It may be that the majority of these searches are by parents whose child has been diagnosed or hospitalized with an RSV infection and are thus looking for information on the infection. Of particular note is the increasing correlation between searches and hospitalizations over time. This could be related to increasing smartphone penetration beginning in 2008 [26], where the parents of hospitalized children are able to search for RSV-related information while sitting in the hospital.

Importantly, of the top ten sites returned in a search for RSV, five were governmental or children’s hospitals, four were medical information websites or Wikipedia, and one was a pharmaceutical company with information on Palivizumab (CDC, lung.org, webmd.org, medlineplus.gov, kidshealth.org, the Mayo Clinic, St Louis Children’s Hospital, Wikipedia, medicinenet.com, and synagis.com, respectively). This indicates that those searching for information on RSV have easy access to evidence-based information and treatment options. These organizations should be com-mended for providing such information, as they are important for clinicians in guiding patients and their families to accurate and detailed information on the typical clinical course of an RSV infection. Clinicians must also be aware of the information-seeking behavior of their patients in order to provide the best care possible.

The observed strong correlation between RSV hospitalizations and internet searches has important implications for the use of search query surveillance as a reliable epidemiological surveillance tool, as well as for subsequent evaluation of situational awareness surrounding RSV hospitalizations. As has been noted, search query surveillance does not suffer from reporting delays inherent to traditional epidemiological reporting. HCUP hospitalization data may take two years from hospitalization to publication of the data; the National Respiratory and Enteric Virus Surveillance System (NREVSS), while providing rolling 3-week averaged weekly tests, it has reporting delays, and potential testing biases in states with small numbers of tests; and the RSVAlert program provides timely RSV test results, but may also may be subject to testing biases. In contrast, Google searches are available immediately and on an hourly basis. This immediacy becomes useful when physicians and public health decision-makers are gauging the current level of RSV transmission and hospitalization rates. Until RSV vaccines are licensed, prophylactic administration of Palivizumab is the only prevention option for the most at-risk children. The work presented here can aid in making the decision to begin this expensive treatment.

Using RSV searches as RSV hospitalization surveillance, we identified two patterns of transmission: first, RSV outbreaks begin in Florida and radiate linearly outward across the country. This has been identified before and may be due to climatological factors and human mobility. Future work can explore this in more detail using metro-level search dynamics. Second, through examining the correlations between the phase-angles and raw time series of searches, we identify three transmission regions: the south and the northeast, the northwest and Appalachia, and the middle of the US. While these three regions are characterized by large metropolitan areas, these transmission regions could be dictated by interstate patterns and population travel and migration. Future work could explore the role of population movement on transmission, perhaps by exploration of historical traffic and/or travel data. We find some differences between clusters identified using phase angles or search volume, notably the inclusion of both coasts in clusters based on search volume. This finding could be due to differential searching behavior in coastal states [27], the clusters identified by phase angles gives a better indicator of transmission clusters as it is based on search timing and not volume. Finally, it remains to be seen how to best implement these findings in a public health setting. Future work in implementation science should explore how these results can better clinical or public health decision-making.

We have shown high correlation between internet searches and RSV hospitalization in the US; it remains an open question as to whether this same correlation will be seen in other countries with differing RSV seasonality, or searching behavior [28]. The dynamics of searching in response to hospital diagnoses may differ between countries and languages. Preliminary results indicate opposite seasonality of RSV searches in Australia corresponding to peak RSV seasons there [29, 30, 31]. Similarly, this work has the limitation that it is potentially influenced by media or shifting search behaviors, both of which adversely affected Google Flu Trends, leading to Google’s discontinuation of that product. Nonetheless, we have shown the face validity of a specific search term relating to a concrete medical outcome.

Finally, as an additional outcome of this work, we have made predictions for numbers of hospitalizations in four states (WA, MD, FL, CT) over three RSV seasons (2016-2018), well ahead of the release of new HCUP, NREVSS, or RSVAlert data. Prospective studies will be able to validate this truly out-of-sample prediction. Here we have linked searches for RSV to hospitalizations for RSV, which suggests that individuals are likely getting accurate medical information when searching for RSV. We have additionally created a website, rsvtrends.org, which will make publically available our RSV hospitalization predictions in near real-time. Our results highlight the utility of searches for medical and public health decision-making, and also highlight the critical need for health professionals to better understand information-seeking because their patients are getting information online.

## Supplementary Table

**Table 1.**
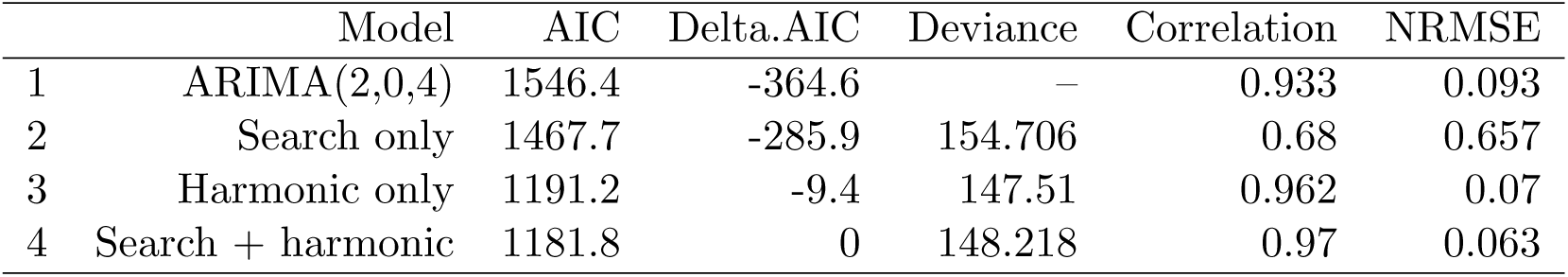
Model performance. Table shows the model performance of the models with and without searches.

|  | Model | AIC | Delta.AIC | Deviance | Correlation | NRMSE |
| --- | --- | --- | --- | --- | --- | --- |
| 1 | ARIMA(2,0,4) | 1546.4 | -364.6 | — | 0.933 | 0.093 |
| 2 | Search only | 1467.7 | -285.9 | 154.706 | 0.68 | 0.657 |
| 3 | Harmonic only | 1191.2 | -9.4 | 147.51 | 0.962 | 0.07 |
| 4 | Search + harmonic | 1181.8 | 0 | 148.218 | 0.97 | 0.063 |

## Supplementary Figures

**Figure 6:**
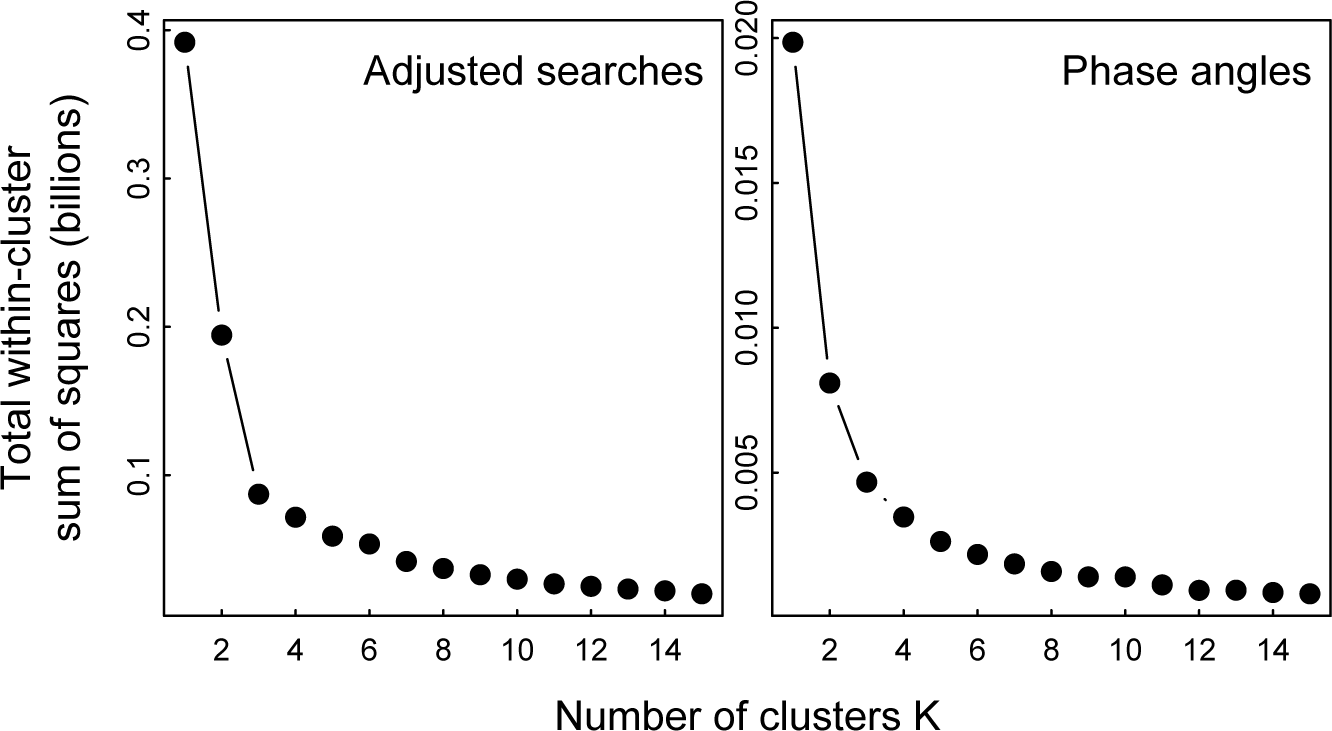
Determination of the optimal number of clusters. Figure shows the total within-cluster sum of squares by cluster size for search volume (left panel) and for phase angles (right panel). Three cluster is the “elobw” of the elbow method where increasing numbers of clusters does not substantially lower the within-cluster sum of squares and thus be the optimal number of clusters.

**Figure 7:**
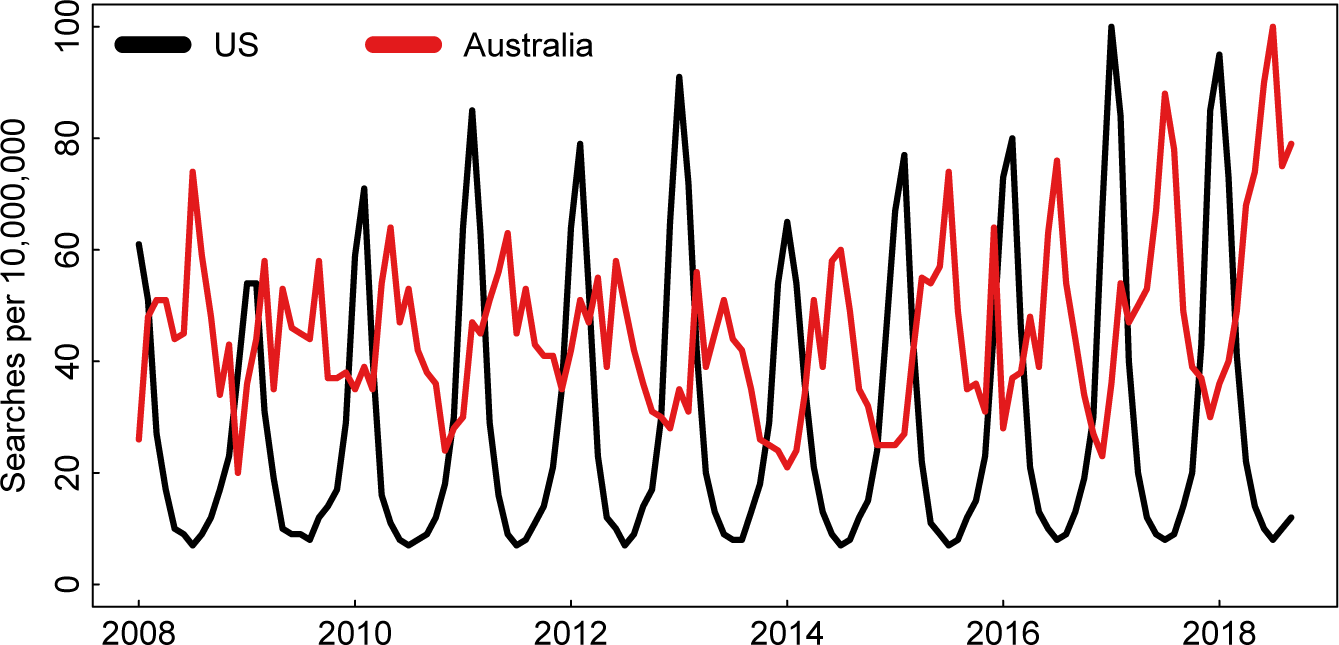
Comparison of RSV search dynamics in the US and Australia. Figure shows the Search volume of RSV per 10,000,000 searches in the US (black) and Australia (red).

## References

[1] B. M. Althouse, S. V. Scarpino, L. A. Meyers, J. W. Ayers, M. Bargsten, J. Baumbach, J. S. Brownstein, L. Castro, H. Clapham, D. A. Cummings, et al., “Enhancing disease surveillance with novel data streams: challenges and opportunities,” EPJ Data Science, vol. 4, no. 1, p. 17, 2015.

[2] J. W. Ayers, B. M. Althouse, and M. Dredze, “Could behavioral medicine lead the web data revolution?,” Jama, vol. 311, no. 14, pp. 1399–1400, 2014.

[3] D. Lazer, R. Kennedy, G. King, and A. Vespignani, “The parable of google flu: traps in big data analysis,” Science, vol. 343, no. 6176, pp. 1203–1205, 2014.

[4] M. Santillana, D. W. Zhang, B. M. Althouse, and J. W. Ayers, “What can digital disease detection learn from (an external revision to) google flu trends?,” American journal of preventive medicine, vol. 47, no. 3, pp. 341–347, 2014.

[5] D. A. Broniatowski, M. J. Paul, and M. Dredze, “National and local influenza surveillance through twitter: an analysis of the 2012-2013 influenza epidemic,” PloS one, vol. 8, no. 12, p. e83672, 2013.

[6] W. W. Thompson, D. K. Shay, E. Weintraub, L. Brammer, N. Cox, L. J. Anderson, and K. Fukuda, “Mortality associated with influenza and respiratory syncytial virus in the united states,” Jama, vol. 289, no. 2, pp. 179–186, 2003.

[7] K. McLaurin, A. Farr, S. Wade, D. Diakun, and D. Stewart, “Respiratory syncytial virus hospitalization outcomes and costs of full-term and preterm infants,” Journal of Perinatology, vol. 36, no. 11, p. 990, 2016.

[8] A. R. Falsey and E. E. Walsh, “Respiratory syncytial virus infection in adults,” Clinical microbiology reviews, vol. 13, no. 3, pp. 371–384, 2000.

[9] PATH, “Rsv vaccine and mab snapshot,” 2017.

[10] I.-R. S. Group et al., “Palivizumab, a humanized respiratory syncytial virus monoclonal antibody, reduces hospitalization from respiratory syncytial virus infection in high-risk infants,” Pediatrics, vol. 102, no. 3, pp. 531–537, 1998.

[11] C. A. Panozzo, L. J. Stockman, A. T. Curns, and L. J. Anderson, “Use of respiratory syncytial virus surveillance data to optimize the timing of immunoprophylaxis,” Pediatrics, pp. peds–2009, 2010.

[12] D. J. Earn, P. Rohani, B. M. Bolker, and B. T. Grenfell, “A simple model for complex dynamical transitions in epidemics,” Science, vol. 287, no. 5453, pp. 667–670, 2000.

[13] E. S. Toner, “Creating situational awareness: A systems approach,” in Medical surge capacity: Workshop summary, National Academies Press, Washington, 2009.

[14] A. Kaji, K. L. Koenig, and T. Bey, “Surge capacity for healthcare systems: a conceptual framework,” Academic Emergency Medicine, vol. 13, no. 11, pp. 1157–1159, 2006.

[15] E. Toner, R. Waldhorn, B. Maldin, L. Borio, J. B. Nuzzo, C. Lam, C. Franco, D. Henderson, T. V. Inglesby, and T. O’Toole, “Hospital preparedness for pandemic influenza,” biosecurity and bioterrorism: biodefense strategy, practice, and science, vol. 4, no. 2, pp. 207–217, 2006.

[16] A. for Healthcare Research and Quality, “Healthcare cost and utilization project (hcup). hcup state inpatient databases (sid) 2004-2015,” 2106.

[17] R. J. Hyndman, Y. Khandakar, et al., Automatic time series for forecasting: the forecast package for R. No. 6/07, Monash University, Department of Econometrics and Business Statistics, 2007.

[18] D. N. Fisman, “Seasonality of infectious diseases,” Annu. Rev. Public Health, vol. 28, pp. 127–143, 2007.

[19] E. Lofgren, N. Fefferman, M. Doshi, and E. N. Naumova, “Assessing seasonal variation in multisource surveillance data: annual harmonic regression,” in Intelligence and Security Informatics: Biosurveillance, pp. 114–123, Springer, 2007.

[20] R Core Team, R: A Language and Environment for Statistical Computing. R Foundation for Statistical Computing, Vienna, Austria, 2018.

[21] B. Cazelles, M. Chavez, D. Berteaux, F. Ménard, J. O. Vik, S. Jenouvrier, and N. C. Stenseth, “Wavelet analysis of ecological time series,” Oecologia, vol. 156, no. 2, pp. 287–304, 2008.

[22] V. E. Pitzer, C. Viboud, W. J. Alonso, T. Wilcox, C. J. Metcalf, C. A. Steiner, A. K. Haynes, and B. T. Grenfell, “Environmental drivers of the spatiotemporal dynamics of respiratory syncytial virus in the united states,” PLoS pathogens, vol. 11, no. 1, p. e1004591, 2015.

[23] R. L. Thorndike, “Who belongs in the family?,” Psychometrika, vol. 18, no. 4, pp. 267–276, 1953.

[24] A. Ng, “Clustering with the k-means algorithm,” Machine Learning, 2012.

[25] D. M. Weinberger, J. L. Warren, C. A. Steiner, V. Charu, C. Viboud, and V. E. Pitzer, “Reduced-dose schedule of prophylaxis based on local data provides near-optimal protection against respiratory syncytial virus,” Clinical Infectious Diseases, vol. 61, no. 4, pp. 506–514, 2015.

[26] A. Lella, “Us smartphone penetration surpassed 80 percent in 2016,” 2017.

[27] J. W. Ayers, B. M. Althouse, J.-P. Allem, E. C. Leas, M. Dredze, and R. S. Williams, “Revisiting the rise of electronic nicotine delivery systems using search query surveillance,” American journal of preventive medicine, vol. 50, no. 6, pp. e173–e181, 2016.

[28] J. W. Ayers, B. M. Althouse, J.-P. Allem, J. N. Rosenquist, and D. E. Ford, “Seasonality in seeking mental health information on google,” American journal of preventive medicine, vol. 44, no. 5, pp. 520–525, 2013.

[29] T. Nolan, C. Borja-Tabora, P. Lopez, L. Weckx, R. Ulloa-Gutierrez, E. Lazcano-Ponce, A. Kerdpanich, M. A. R. Weber, A. Mascareñas de Los Santos, J.-C. Tinoco, et al., “Prevalence and incidence of respiratory syncytial virus and other respiratory viral infections in children aged 6 months to 10 years with influenza-like illness enrolled in a randomized trial,” Clinical Infectious Diseases, vol. 60, no. 11, pp. e80–e89, 2015.

[30] A. Dede, D. Isaacs, P. J. Torzillo, J. Wakerman, R. Roseby, R. Fahy, T. Clothier, A. White, and P. Kitto, “Respiratory syncytial virus infections in central australia,” Journal of paediatrics and child health, vol. 46, no. 1-2, pp. 35–39, 2010.

[31] J. Druce, T. Tran, H. Kelly, M. Kaye, D. Chibo, R. Kostecki, A. Amiri, M. Catton, and C. Birch, “Laboratory diagnosis and surveillance of human respiratory viruses by pcr in victoria, australia, 2002– 2003,” Journal of medical virology, vol. 75, no. 1, pp. 122–129, 2005.

